# Lactate dehydrogenases amplify reactive oxygen species in cancer cells in response to oxidative stimuli

**DOI:** 10.1101/376210

**Authors:** Hao Wu, Yuqi Wang, Mingfeng Ying, Xun Hu

## Abstract

Previous studies have revealed that lactate dehydrogenase A (LDHA) exhibited an indirect antioxidative activity, which played a crucial role in preventing cancer cells from oxidative stress. Here we demonstrated that, apart from antioxidative activities, LDHA and LDHB displayed prooxidative activity in cancer cells. In aqueous phase, LDHA and B both exhibited ROS-generating activity and LDHB was more active than LDHA. In cancer cells, the dominant antioxidative activity of LDH can be switched to dominant prooxidative activity or vice versa, depending on the strength of oxidative stimuli, indicating that LDH were bifunctional. Moreover, we demonstrated that mitochondrial superoxide served as an initiator to trigger LDH-catalyzed amplification of ROS. The oxidative stimuli, such as modulators of electron transfer chain and anticancer agents that kill cancer cells via ROS induction, induced a ROS generation involving 2 phases, induction of mitochondrial superoxide and amplification of ROS by LDH.

## Introduction

In cells, superoxide is mainly produced in mitochondrion, in which electron leaks from the electron transfer chain (ETC) in the inner membrane of mitochondria and is captured by molecular oxygen. It is estimated that about 3% of the molecular oxygen reduced in the mitochondria produce superoxide(***Freeman and Crapo,1981; Park, et al.,1998; St-Pierre, et al.,2002***). The other sources of superoxide generation in cells include NAPDH oxidase, xanthine oxidase, cytochrome P450 peroxidases, indoleamine 2,3-dioxygenase, tryptophan dioxygenase(***Fridovich, 1995; Imlay,2002; Satoshi Aoki,1989***). When tetrahydobiopterin and arginine is low, nitric oxide synthase also generate superoxide(***Berka, et al.,2004; Xia, et al.,1998***). It is recognized that molecules in cells are initially slowly oxidized, but once superoxide is generated and accumulated, it sets up a chain of free radical reactions and accelerates oxidation of molecules in cells.

One key question is if some enzymes per se do not generate superoxide but could amplify the free radical reactions. Lactate dehydrogenase is a candidate. Previously, Chan and Bielski reported that in aqueous phase, superoxide and hydroperoxyl radicals are in equilibrium with a pK = 4.8, and this amount of superoxide is sufficient to initiate and amplify a chain of free radical reaction on LDH-bound NADH, as illustrated by the following chemical reactions(***Bielski and Chan,1976; Chan and Bielski,1974***).

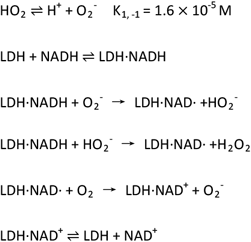

As a result, in each chain of reactions, a molecule of superoxide and a molecule of hydrogen peroxide are generated. This established the theoretical basis for LDH to serve as a universal amplifier of reactive oxygen species.

LDH is abundant and mitochondria are functional in most cancer cells. It is possible that superoxide generated from the mitochondria or other sources could initiate LDH to amplify the chain of free radical reactions in cancer cells.

## Results and Discussion

### 1. ROS-generation activity of LDH A and B

In the reaction system, ROS generation was dependent on the concentration of human LDHA and B (Figure 1a) and the concentration of NADH (Figure 1b). ROS generation mediated by LDHA and LDHB was inhibited by SOD (Figure 1c). The signal saturated at high enzyme concentration is probably due to the limited concentration of superoxide in the reaction system (Figure 1a). These experiments reproduced the study by Chan and Bielski, who discovered that the LDH catalyzed a chain of free radical reactions, which is initiated by superoxide in an aqueous solution, in which superoxide and hydroperoxyl radicals are in equilibrium with a pK = 4.8(***Chan and Bielski,1974***). Moreover, the ROS generation was inhibited by LDH inhibitor FX11 (NAD binding site) (Figure 1d) and oxamate (pyruvate binding site) (Figure 1e), with the former being a significantly stronger inhibitor than the latter. The bovine LDHA and LDHB showed the same pattern (Figure 1f – j).

**Figure 1.**
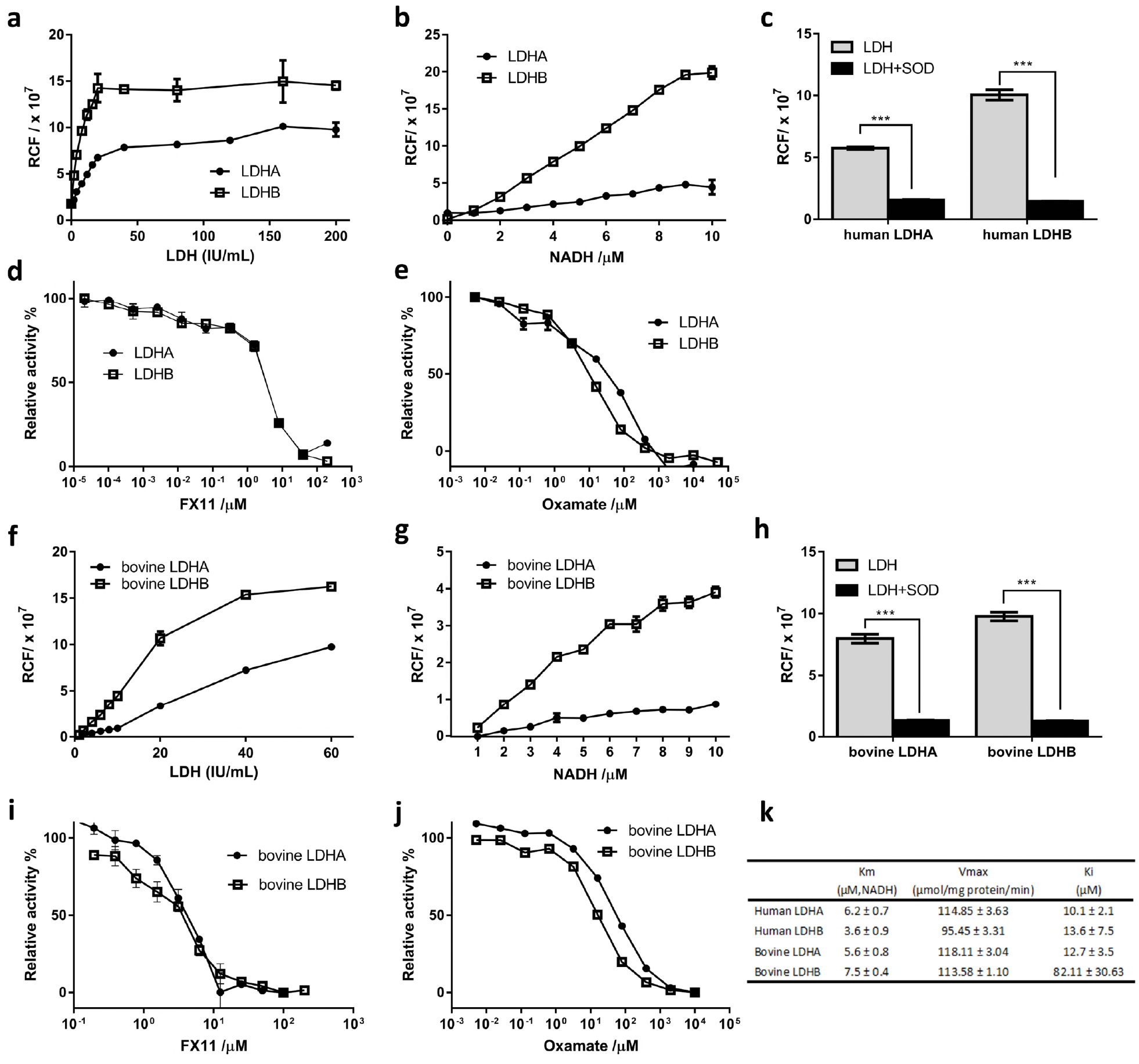
The ROS-generating activity of LDHA and B. (a-e human LDH, f–j bovine LDH) (a & f) LDH activity dependent ROS generation. 1 unit of LDH refers the conversion of 1 μmole of pyruvate into lactate in 1 minute. (b & g) NADH dependent ROS generation by LDH. (c & h) ROS-generating activity of LDH with or without SOD. (d, e, i, j) LDH inhibitors FX11 and oxamate inhibit LDH-catalyzed ROS generation. (k) The kinetic parameters of human and bovine LDH in converting pyruvate to lactate and Ki refers the LDH inhibitor FX11. Experiments were repeated 3 times and one representative data is showed and expressed as mean ± SD. n = 3. The experimental details are described in Materials and Methods.

An interesting finding was that LDHB per unit wise displayed a significantly higher activity than LDHA with respect to ROS generating (Figure 1a, 1b, 1f, 1g). LDHA and LDHB showed similar activities in terms of pyruvate to lactate conversion (Figure 1k)

### 2. Defining the antioxidative and prooxidative activities of LDH in cells

Previous reports demonstrated that inhibition of LDH by FX11 or LDH knockdown resulted in a significant increase of cellular ROS (***Arseneault, et al.,2013; Le A, et al.,2010; Newington, et al.,2012; Seth, et al.,2011; Xie, et al.,2014***). Presumably, inhibition of LDH diverts pyruvate for complete oxidation in mitochondria and such diversion may significantly increase electron flux rate on the ETC. As the electron leakage is proportional to the electron flux rate along ETC, LDH inhibition eventually resulted in an increase of mitochondrial superoxide, leading to oxidative stress.

We also reproduced their results, i.e., inhibition of LDH by FX11 or LDH knockdown in Hela cells resulted in a significant increase of cellular ROS (Figure 2a & b). FX11 inhibition apparently exerted a stronger effect on ROS generation than LDH knockdown, because FX11 can inhibit LDH activity more thoroughly than LDH knockdown.

**Figure 2.**
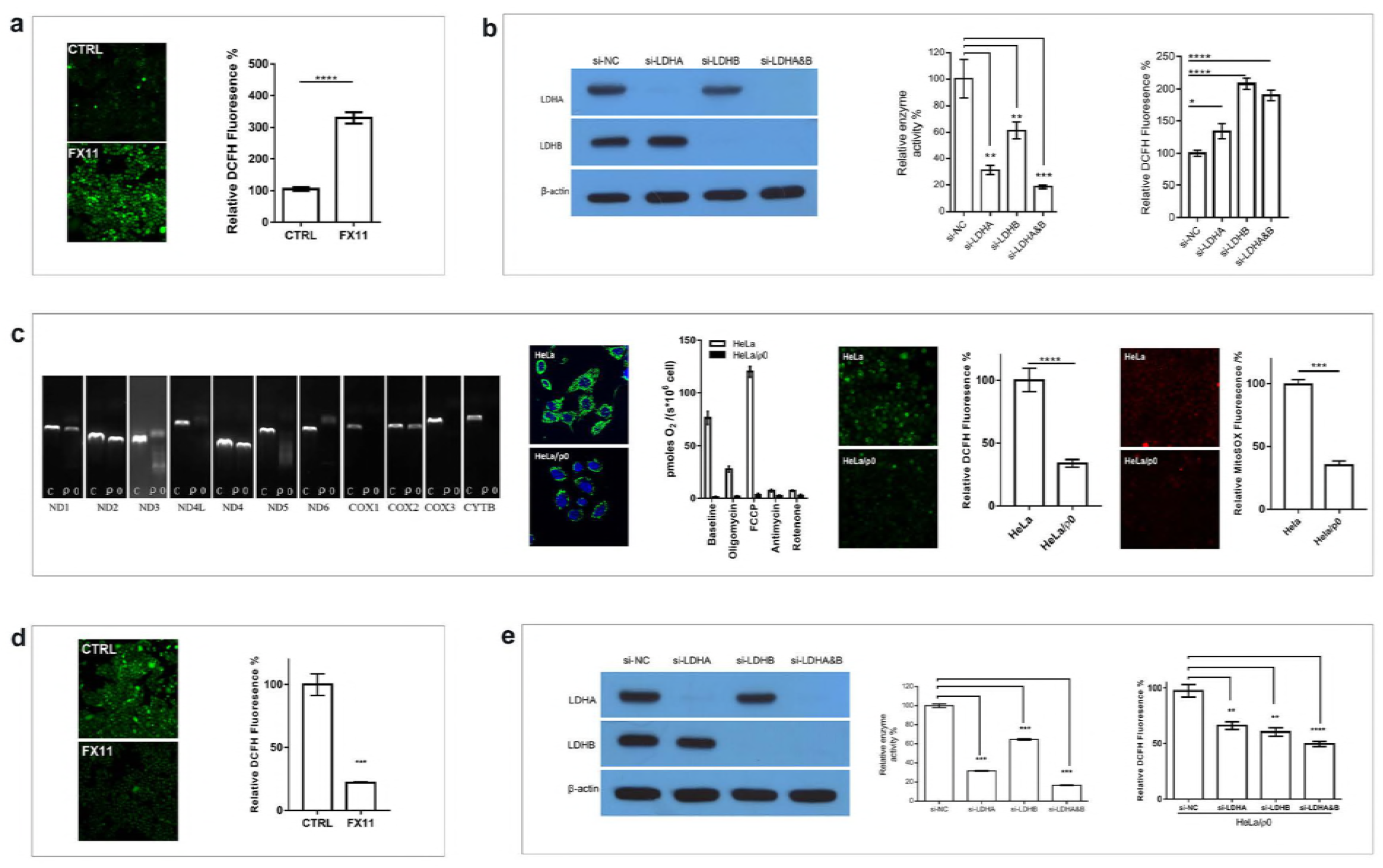
The antioxidative activity of LDH in Hela and prooxidant activity of LDH in Hela/p0 cells (a & b, Hela cells, c – e Hela/p0 cells). (a) LDH inhibitor FX11 enhances ROS production (DCFH). (b) LDH A or B or both knockdown enhances ROS production (DCFH). From left to right, Western blot of LDH; relative LDH activity assayed by converting pyruvate to lactate; relative ROS levels in cells. (c) characterization of Hela/p0. from left to right: mitochondrial DNA electrophoresis of Hela and HeLa/p0 cells; MitoTraker - a Mitochondrion-selective probe stain of Hela and Hela/p0 cells; oxygen consumption rate in comparison to Hela cells; DCFH signal in comparison to Hela cells; MitoSOX™ Red signal in comparison to Hela cells. (d) FX11 reduces DCFH signal in Hela/p0 cells. (e) LDHA or LDHB or both knockdown reduces DCFH signals in Hela/p0 cells. From left to right, Western blot of LDH; relative LDH activity assayed by converting pyruvate to lactate; relative DCFH signals in cells. Data were confirmed by at least 2 independent experiments. ND1: MT-ND1, mitochondrially encoded NADH dehydrogenase 1; ND2: MT-ND2, mitochondrially encoded NADH dehydrogenase 2; ND3: MT-ND3, mitochondrially encoded NADH dehydrogenase 3; ND4L: MT-ND4L, mitochondrially encoded NADH 4L dehydrogenase; ND4: MT-ND4, mitochondrially encoded NADH dehydrogenase 4; ND5: MT-ND5, mitochondrially encoded NADH dehydrogenase 5; ND6: MT-ND6, mitochondrially encoded NADH dehydrogenase 6; COX1: MT-CO1, mitochondrially encoded cytochrome c oxidase I; COX1: MT-CO1, mitochondrially encoded cytochrome c oxidase II; COX3: MT-CO3, mitochondrially encoded cytochrome c oxidase III; CYTB: MT-CYB, mitochondrially encoded cytochrome b. The experimental details are described in Materials and Methods.

We then tested the effect of inhibiting LDH in Hela/ρ0 cells on ROS production. As Hela/ρ0 cells are mitochondrial DNA deficient and lack some essential components of ETC complex (Figure 2c), inhibition of LDH in these cells should not lead to an enhanced electron flux on ETC hence should not increase ROS, instead, if LDH could produce ROS, the inhibition of LDH would reduce cellular ROS. The basal levels of mitochondrial superoxide and total cellular ROS in HeLa/ρ0 cells were significantly lower than those in Hela cells (Figure 2c). FX11 treatment resulted in a significant decrease of cellular ROS (Figure 2d). LDHA, or LDHB or both Knockdown also led to a reduction of cellular ROS in HeLa/ρ0 cells (Figure 2e).

Taken together, we demonstrated that LDHA and B could also generate ROS in mitochondrial DNA-deficient ρ0 cells and in addition the results were consistent with the previous reports in terms of antioxidative activity of LDHA (***Arseneault, et al.,2013; Le A, et al.,2010; Newington, et al.,2012; Seth, et al.,2011; Xie, et al.,2014***). Thus, LDH is bifunctional, indirect antioxidative activity proposed by other investigators (***Arseneault, et al.,2013; Le A, et al.,2010; Newington, et al.,2012; Seth, et al.,2011; Xie, et al.,2014***) and direct prooxidative activity demonstrated in ρ0 cells by us.

### 3. The relationship between induction of mitochondrial superoxide and LDH-mediated amplification of cellular ROS

According to the reaction mechanism, superoxide is so far the strongest initiator to trigger LDH-catalyzed free radical reaction(***Bielski and Chan,1976; Chan and Bielski,1974; Petrat, et al.,2005***). Mitochondrion is the major intracellular site to generate superoxide. We assume that there was an interaction between mitochondrial superoxide and total cellular ROS. Superoxide generated from mitochondria may serve as initiator to trigger LDH-catalyzed free radical reaction, which amplifies cellular ROS.

We treated cells with rotenone, antimycin, FCCP, or oligomycin. Rotenone inhibits electron flow in complex I and the inhibition would saturate complex I with electron, leading to electron leakage and producing superoxide; antimycin stops electron flux at complex III, leading to electron saturation at complex I, complex II, ubiquinone, and complex III, eventually increasing electron leakage(***Chen, et al.,2003; Li, et al.,2003; Wardman,2007***). FCCP, an uncoupler of oxidative phosphorylation, maximizes the electron flux rate along ETC, thus also increases probability of electron leakage. Oligomycin is an inhibitor of F_o_ part of ATP synthase. In theory, these agents would significantly enhance superoxide production in mitochondria(***Turrens,2003***).

Throughout the text, unless otherwise stated, mitochondrial superoxide refers the fluorescent intensity of MitoSOX™ Red and total cellular ROS refers the fluorescent intensity of DCFH.

These agents increased mitochondrial superoxide and total cellular ROS (Figure 3a, upper and middle panels). In the presence of FX11, total cellular ROS level with or without treatment of rotenone, antimycin, FCCP, or oligomycin was comparable with each other, i.e., FX11 increased cellular ROS in the control cells but reduced ROS in the treated cells to a similar level (Figure 3a, down panels). The results suggest the double-edged function of LDH: without treatment of rotenone, antimycin, FCCP, or oligomycin, the indirect antioxidative activity of LDH was dominant, agreeable with previous reports(***Arseneault, et al.,2013; Le A, et al.,2010; Newington, et al.,2012; Seth, et al.,2011; Xie, et al.,2014***), while with treatment of these agents, the prooxidative activity of LDH became dominant.

**Figure 3.**
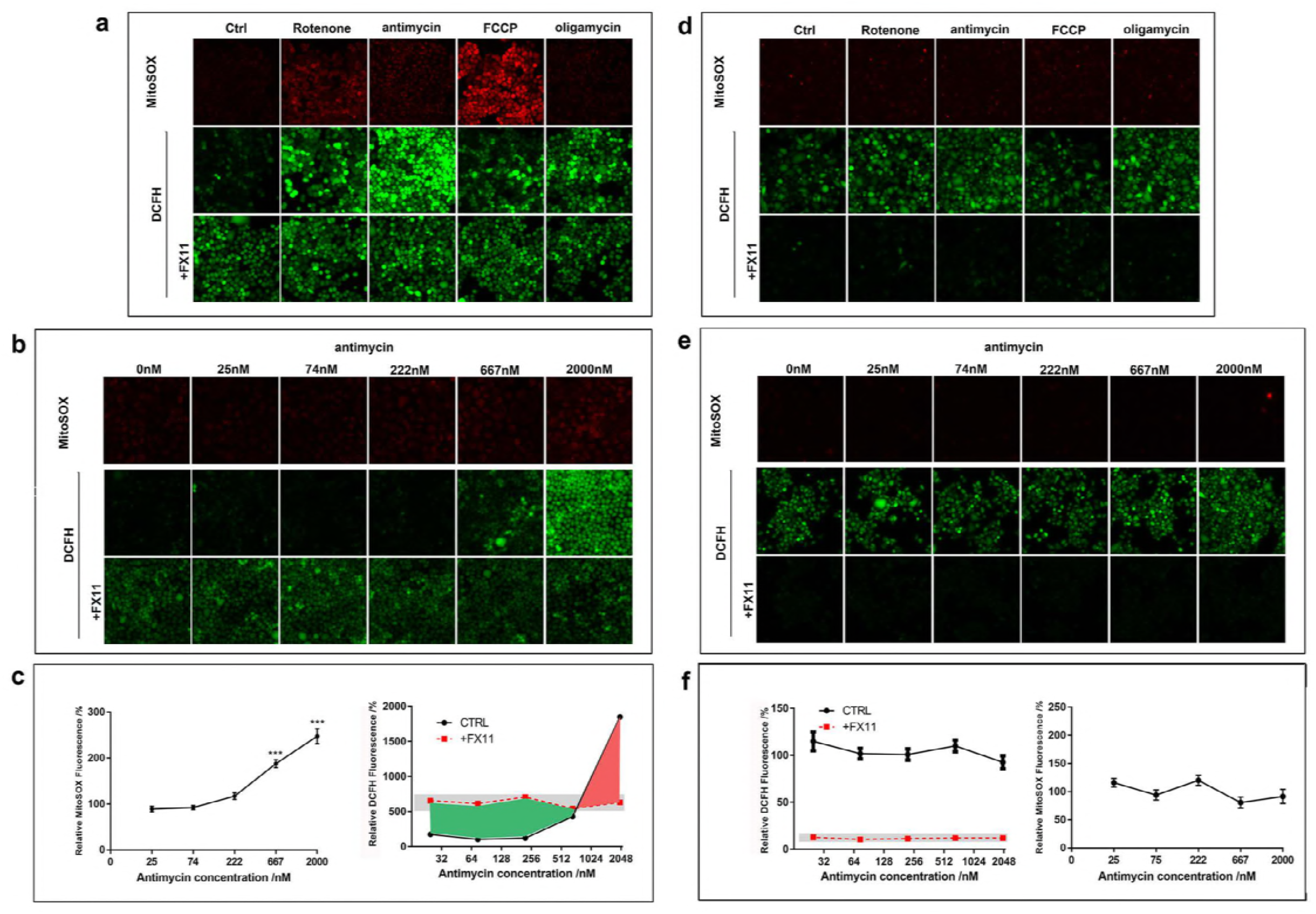
Revelation of the dominant antioxidative and dominant prooxidative activity of LDH in Hela and Hela/ρ0 cells. (a-c, Hela cells; d-f, Hela/ρ0 cells). (a & d) Rotenone-, antimycin-, FCCP-, or oligomycin-induced production of mitochondrial superoxide (MitoSOX™ Red) and total cellular ROS (DCFH) in Hela and Hela/ρ0 cells with or without FX11. (b & e) Antimycin showed concentration-dependent induction of mitochondrial superoxide(MitoSOX™ Red) and total cellular ROS (DCFH)in Hela and Hela/ρ0 cells with or without FX11. (c & f) The statistical data of the b & e. The green and red area represent the conversion from dominant antioxidative to dominant prooxidant activity of LDH or vice versa. The intersection point represents equal antioxidative and prooxidant activity of LDH. Data were confirmed by at least 2 independent experiments. The experimental details are described in Materials and Methods.

The above results suggested that the amount of superoxide generated from mitochondria was a key to regulate LDH between the indirect antioxidative activity and direct prooxidative activity in cells. To further investigate this, we treated cells with a serial concentrations of antimycin and observed the following phenomenon: (1) without FX11, antimycin induced an increase of mitochondrial superoxide and cellular ROS (figure 3b upper and middle panels); (2) with or without antimycin, FX11 brought the cellular ROS to a similar level (figure 3b down panel); (3) There is an intersection point of 2 lines (figure 3c), the green area indicates the dominant indirect antioxidative activities and the red area reflects the dominant direct prooxidative activities of LDH, the intersection point reflects the equal anti- and pro-oxidative activities of LDH under the experimental setting (figure 3c).

Then, we used Hela/ρ0 cells to perform the same experiment. Since ETC was defective, these agents had no effect on mitochondrial superoxide, neither total cellular ROS (Figure 3d & e). In the presence of FX11, cellular ROS reduced significantly (Figure 3f).

Taken together, the above experiments demonstrated a link between mitochondrial superoxide and total cellular ROS, suggesting mitochondrial superoxide as the initiator to activate LDH-catalyzed free radical reaction, that amplifies total cellular ROS in cells. In addition, the relationship between the indirect antioxidative activity of and the direct prooxidative activity of LDH was also captured (Figure 3c).

### 4. LDHA or LDHB knockout do not affect mitochondrial superoxide but decrease cellular ROS induced by the ETC modulators

We sought to use LDH knockout cells (Figure 4a, Supplementary figure 1a) to study the relationship between LDH activity and cellular ROS levels. The subtype of LDH in Hela/LDHA_KO_ and Hela/LDHB_KO_ cells was LDHB and LDHA, respectively. Although LDH activity of Hela/LDHA_KO_ and Hela/LDHB_KO_ cells assayed by pyruvate and lactate conversion differed by about 2 folds (Figure 4a, the second panel from the left), in terms of ROS generating, Hela/LDHA_KO_ and Hela/LDHB_KO_ cells were close (Figure 4a, the third panel from the left), as LDHB was about 1.6 folds more potent than LDHA in generating ROS (Figure 1a & b).

**Figure 4.**
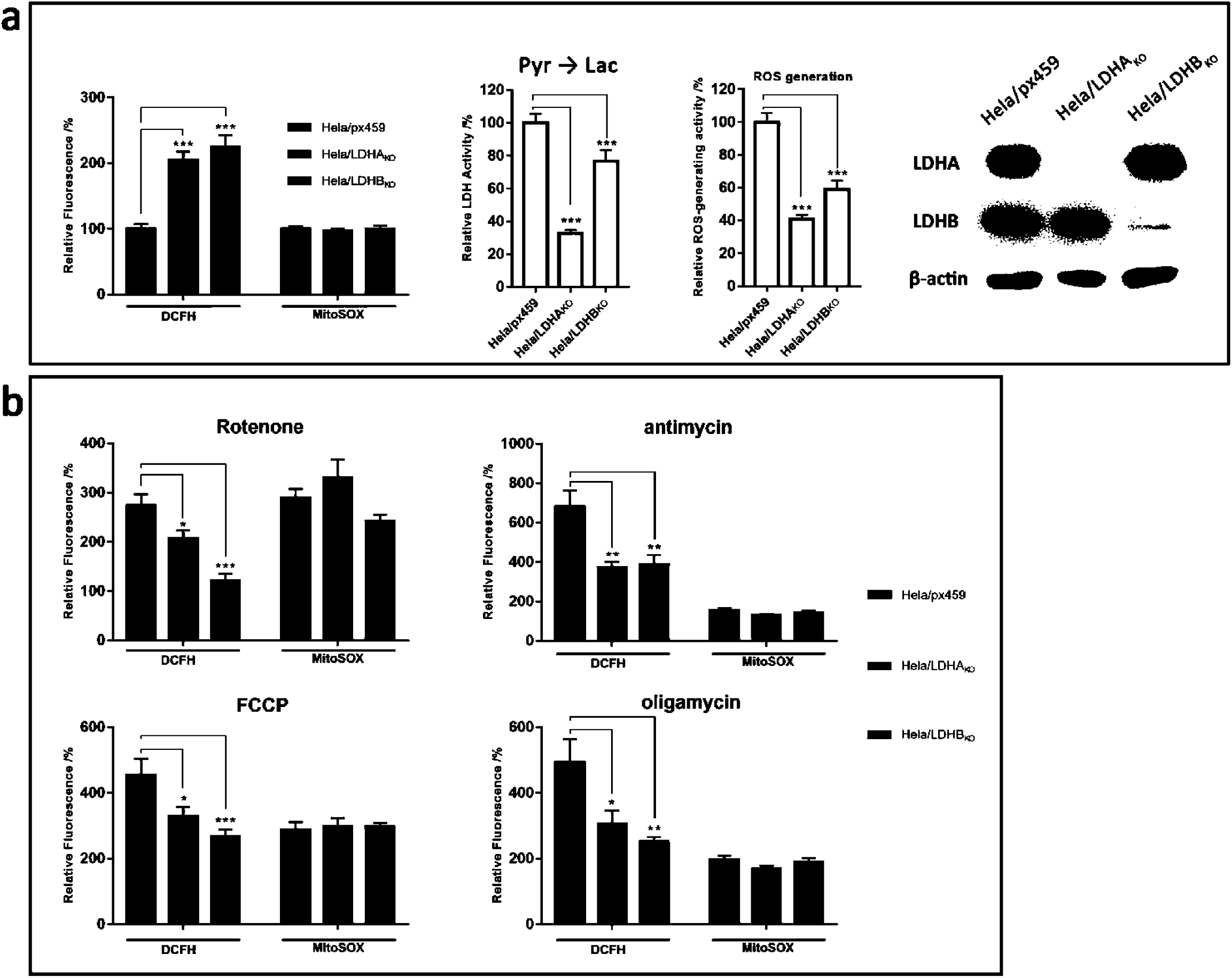
The relationship between cellular ROS and LDH knockout. (a) Characterization of Hela/LDHA_KO_, and Hela/LDHB_KO_ cells. From left to right: relative DCFH and MitoSOX™ Red signals; LDH activity assayed by converting pyruvate to lactate; the ROS generation activity calculated according to figure 1a, in which LDHB is about 1.6 folds as much as LDHA; Western blot of LDH. (b) rotenone-, antimycin-, FCCP-, or oligomycin-induced production of mitochondrial superoxide (MitoSOX™ Red) and total cellular ROS (DCFH). The fluorescence intensity is relative to untreated Hela/px459 cells, which is 100%. Data were confirmed by at least 2 independent experiments. The experimental details are described in Materials and Methods.

The empty vector control cell line Hela/px459, and LDH knockout cell lines Hela/LDHA_KO_ and Hela/LDHB_KO_ cells displayed similar mitochondrial superoxide levels, indicating that LDHA or LDHB knockout did not affect mitochondrial superoxide (Figure 4a, the first panel from the left). This is surprising, as we expected that LDHA or LDHB knockout should increase mitochondrial superoxide, according to the hypothesis (***Arseneault, et al.,2013; Le A, et al.,2010; Newington, et al.,2012; Seth, et al.,2011; Xie, et al.,2014***). However, as this is not the focus of this study, we did not further pursue the molecular mechanism. Hela/LDHA_KO_ and Hela/LDHB_KO_ cells both displayed higher cellular ROS level than Hela (Figure 4a, the first panel from the left), consistent with the data in Figure 2b and the previous report(***Le A, et al.,2010***).

Rotenone, antimycin, FCCP, and oligomycin all significantly increased mitochondrial superoxide in Hela/px459, Hela/LDHA_KO_ and Hela/LDHB_KO_ cells, whose levels were comparable with each other (Figure 4b). On the other hand, these agents induced a significantly higher level of cellular ROS in control cells than in Hela/LDHA_KO_ and Hela/LDHB_KO_ cells (Figure 4b), indicating that cellular ROS level was positively associated with LDH activity. We then used the empty vector control cell line 4T1/px459, and LDH knockout cell lines 4T1/LDHA_KO_, 4T1/LDHB_KO_ to repeat above experiment and obtained the similar results (Supplementary figure 1b, Supplementary figure 2).

Taken together, ETC modulators enhanced production of mitochondrial superoxide and cellular ROS, LDHA or B knockout selectively inhibited cellular ROS production induced by these agents without affecting mitochondrial superoxide production induced by these agents, indicating that LDHA and B were only responsible for cellular ROS amplification.

### 5. **LDH amplifies ROS induced by anticancer agents**

ROS-based anticancer therapy is emerging (***Trachootham, et al.,2009***). Although PL (***Raj, et al.,2011***) and PEITC(***Trachootham, et al.,2006***) are typical anticancer agents that kill cancer cells through induction of ROS, the underlying molecular mechanism by which they induce ROS is largely unknown.

We proposed that PL- or PEITC-induced ROS production is composed of 2 phases, initiation and amplification. In order to delineate this, we used a pair of cells, Hela and its ρ0 cells (Figure 5a). In Hela cells, PL or PEITC induced a significant elevation of mitochondrial superoxide and total cellular ROS. On the other hand, PL or PEITC did not affect mitochondrial superoxide nor total cellular ROS in Hela/ρ0 cells. We then used another pair of cells, HCT116 and HCT116/ρ0 cells (Figure 5b and Supplementary figure 3), and obtained the same results. The results suggest that, similar to antimycin (Figure 3), PL and PEITC depend on mitochondria for the induction of cellular ROS.

**Figure 5.**
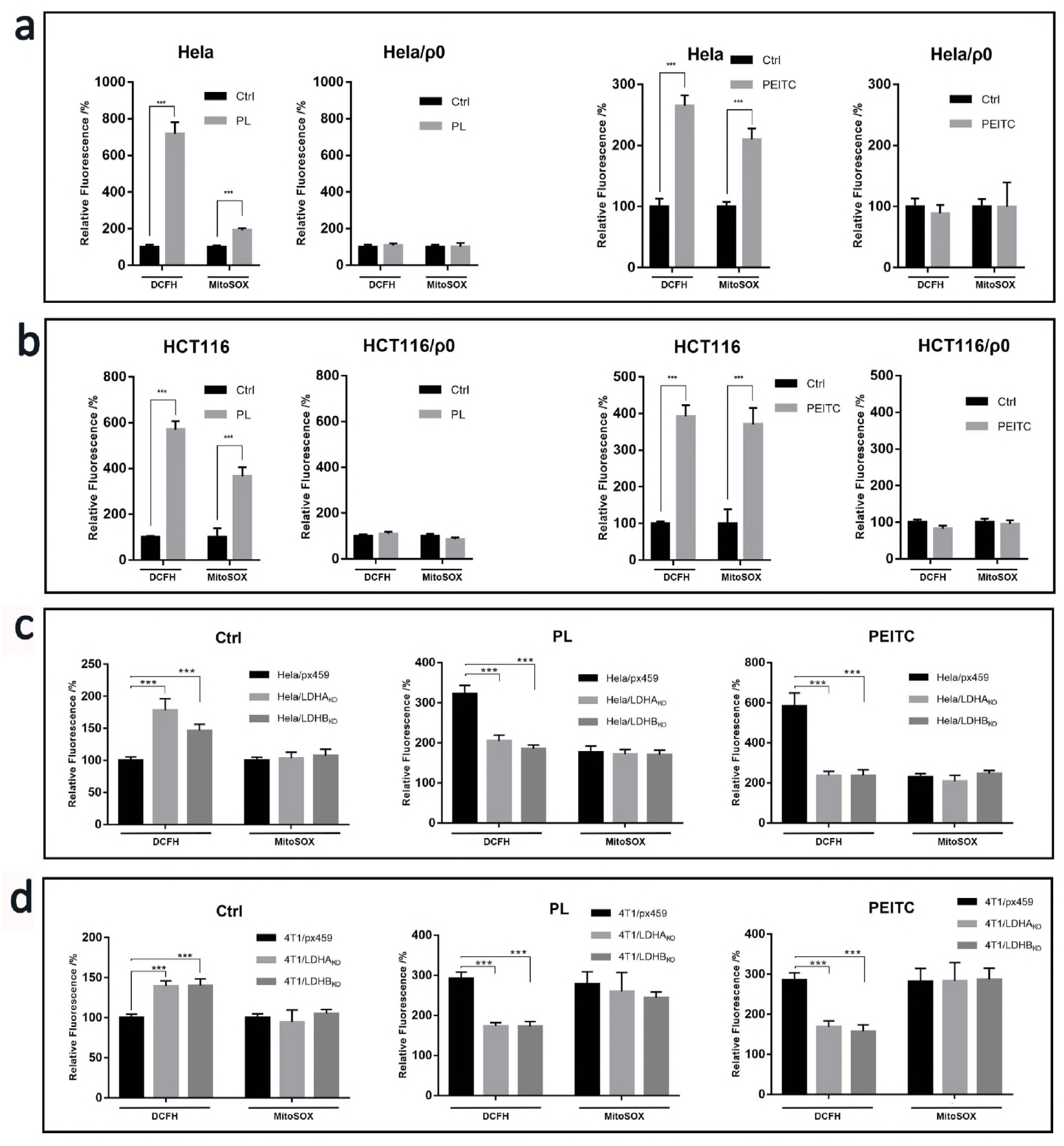
The relationship between PL- or PEITC-induced generation of ROS and LDH. (a & b) PL- or PEITC-induced mitochondrial superoxide (MitoSOX™ Red) and total cellular ROS (DCFH) in Hela and Hela/ρ0 cells, and in HCT116 and HCT116/ ρ0 cells. (c) PL- or PEITC-induced mitochondrial superoxide and cellular ROS in empty vector control Hela/px459 cell, and LDH knockout Hela/LDHA_KO_, and Hela/LDHB_KO_ cells. Ctrl, cells not treated by PL or PEITC. In panels of PL and PEITIC treatment, the fluorescence is relative to untreated Hela cells, which is 100%. (d) PL- or PEITC-induced mitochondrial superoxide and cellular ROS in empty vector control cell line 4T1/px459, and LDH knockout 4T1/LDHA_KO_, and 4T1/LDHB_KO_ cells. Ctrl, cells not treated by PL or PEITC. In panels of PL and PEITIC treatment, the fluorescence intensity is relative to untreated Hela/px459 cells, which is 100%. Data were confirmed by at least 2 independent experiments. The experimental details are described in Materials and Methods.

We then treated Hela/px459 cells, Hela/ LDHA_KO_, and Hela/ LDHA_KO_ cells with PL or PEITC. While Hela/px459 cells showed a significant increase of cellular ROS and mitochondrial superoxide, Hela/ LDHA_KO_, and Hela/ LDHB_KO_ exhibited an increase of mitochondrial superoxide comparable to Hela/px459 cells but a moderate or not significant increase of cellular ROS (Figure 5c). We then treated 4T1/px459, 4T1/ LDHA_KO_, and 4T1/ LDHB_KO_ cells with PL or PEITC, and obtained the same results (Figure 5d). The results demonstrated that LDH knockout selectively inhibited total cellular ROS induced by PL or PEITC without affecting mitochondrial superoxide induced by PL or PEITC, indicating that LDHA and B were only responsible for cellular ROS amplification.

Doxorubicin enhances ROS production through generating superoxide via quinone one-electron redox cycling (***Gewirtz,1999***). We tested if LDH was involved in the amplification of doxorubicin-induced ROS. We treated wild type cells (Hela/px459 and 4T1/px459) and LDHA or LDHB knockout cells with the drug and observed a significant reduction of cellular ROS in LDHA or LDHB knockout cells (Figure 6a). The results supported that LDH was involved in the amplification of doxorubicin-induced ROS.

**Figure 6.**
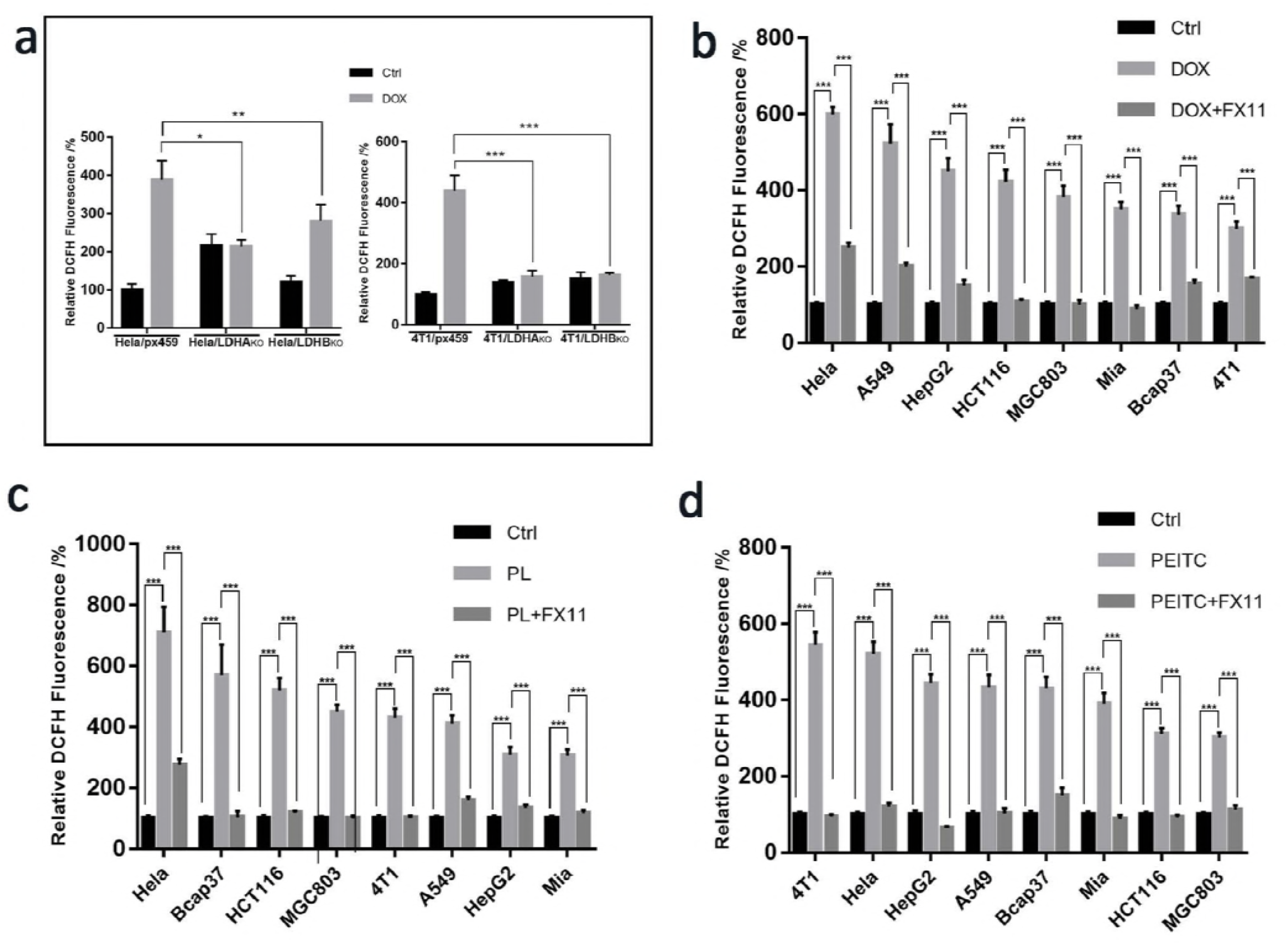
The relationship between cellular ROS levels and activity of LDH. (a) Doxorubicin-induced ROS (DCFH) in Hela/px459, Hela/LDHA_KO_, and Hela/LDHB_KO_ cells, and in 4T1/px459, 4T1/LDHA_KO_, and 4T1/LDHB_KO_ cells. (b - d) Doxorubicin-, PL-, or PEITC-induced ROS (DCFH) in 8 cell lines with or without FX11. Data were confirmed by at least 2 independent experiments. The experimental details are described in Materials and Methods.

In order to confirm if LDH amplification of ROS induced by PL, PEITC, or DOX is a general phenomenon in cancer cells, we treated 8 cell lines (Figure 6b, c & d) with these drugs with or without FX11. The results showed that FX11 could abolish ROS production induced by these drugs.

### 6. **A proposed mechanism of ROS initiation and amplification**

Chan and Bielski (***Bielski and Chan,1976; Chan and Bielski,1974***) established the theoretical basis by which superoxide triggers LDH to catalyze a free radical chain reaction. Molecular oxygen, hydrogen peroxide, peroxynitrite, superoxide all could initiate a chain of free radical reactions on LDH-bounded NADH(***Petrat, et al.,2005***). Thus, in principle, LDH could be a universal ROS amplifier in cells. Experimentally, we provide evidence that LDH is responsible for the amplification of ROS induced by rotenone, antimycin, FCCP, oligomycin, PL, PEITC, and doxorubicin. On the above theoretical and experimental basis, we propose a mechanism by which LDH amplify cellular ROS (Figure 7). Oxidative stimuli enhance electron leakage from ETC in the inner membrane of mitochondrion and the leaked electron is captured by molecular oxygen to form superoxide, which then initiates free radical chain reaction on LDH-bound NADH, ultimately leading to amplification of cellular ROS. Alternatively, superoxide generated from other sources, such as superoxide produced through quinone one-electron redox cycling, could also initiate this amplification. The high glycolysis rate in cancer cells continuously provides glyceraldehyde 3-phosphate, which is a major source of hydride donor that reduces NAD to NADH catalyzed by GAPDH.

**Figure 7.**
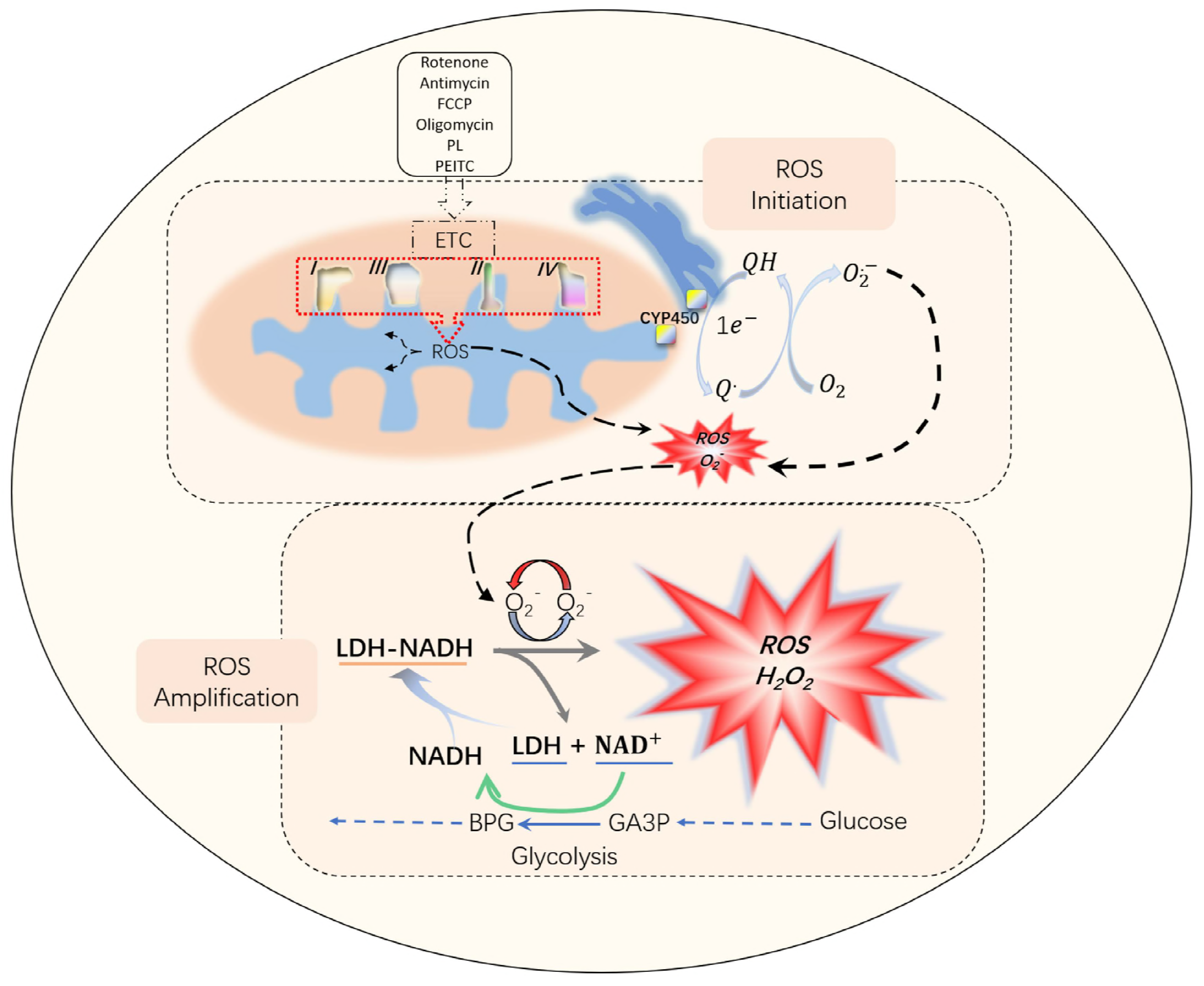
LDH is involved in ROS amplification in cancer cells in response to oxidative stimuli. Oxidative stimuli (rotenone, antimycin, oligomycin, FCCP, PL, PEITC) cause an increase of electron leakage from ETC in the inner membrane of mitochondria and the electron is captured by molecular oxygen to produce superoxide, which serves as an initiator to trigger a free radical chain reaction on LDH-bound NADH, leading to amplification of total cellular ROS. Alternatively, other sources of superoxide, such as superoxide produced from quinone one-electron redox cycling catalyzed by CYP450 can also serve as an initiator to trigger LDH-mediated ROS amplification. Glycolysis continuously provides hydride to reduce NAD+ to NADH at the step of GAPDH. FCCP, carbonyl cyanide 4-(trifluoromethoxy)phenylhydrazone; PL, piperlongumine; PEITC, β -phenylethyl isothiocyanate. GA3P, glyceraldehyde-3 phosphate; BPG, 1,3-bisphosphateglyerate; GAPDH, glyceraldehyde-3 phosphate dehydrogenase.

As the last enzyme of glycolysis, LDH-A plays a crucial role in Warburg effect of cancer cells. It is well recognized that LDH-A is an emerging target to inhibit cancerous glycolysis, tumorigenesis and tumor progression(***Le A, et al.,2010***). The lactate produced by LDH can also be used as nutrient, which is primarily reduced to pyruvate by LDH, to support the growth of other cancer cells or tumor stromal cells and promote tumor progression and metastasis further. Some small inhibitors of LDH-A have been tested in pre-clinical models(***van der Mijn, et al.,2016***). Our data suggest that LDH could serve as an amplifier of ROS. Based on the cellular condition, it could show either overall anti-oxidant or pro-oxidant activity. Therefore, LDH is a cross point of metabolism and ROS regulation. This suggests that more care should be taken in the future designing of therapeutic methods targeting LDH or lactate metabolism.

## Materials and Methods

### Cancer cell lines

Human cervical cancer Hela cells, human breast cancer Bcap37 cells, mouse breast cancer 4T1 cells, human gastric cancer MGC803 cells were maintained in complete RPMI-1640 medium (Gibco) with 10% FBS and 2 mM L-glutamine. Human colorectal carcinoma HCT116 were maintained in complete MyCoy’s 5A medium (Gibco) with 10% FBS and 2 mM L-glutamine. Human liver cancer HepG2 cells were maintained in complete MEM medium (Gibco) with 10% FBS and 2 mM L-glutamine. Human lung cancer A549 cells were maintained in complete F-12K medium (Gibco) with 10% FBS and 2 mM L-glutamine. Human pancreatic cancer Mia cells were maintained in complete DMEM medium (Gibco) with 10% FBS and 2 mM L-glutamine. The ρ^0^ cells were maintained in corresponding complete medium with 50 ng/mL Ethidium bromide (EB), 2 mM pyruvate, and 50 μg/mL uridine. All the medium contained 100 U/ml penicillin/streptomycin.

### Mitochondrial DNA-depleted ρ _0_ cell induction and characterization

HeLa cell and HCT116 cell were seeded into 6-well plate at density of 100 cells /well. 50 ng/mL Ethidium bromide (EB), 2 mM pyruvate, 50 μg/mL uridine were added to culture medium after overnight incubation. The cells were sub-cultured in the same condition when necessary. One and half mouths later, cells were collected, and total DNA was extracted by Blood & Cell Culture DNA Mini Kit (Qiagen). The mitochondrial DNA encoded genes were amplified with the following primers: ND1-F:

5’-CAACATCGAATACGCCGCAG-3’, ND1-R:
5’-AATCGGGGGTATGCTGTTCG-3’, ND2-F:
5’-AACCCTCGTTCCACAGAAGC-3’, ND2-R:
5’-AGCTTGTTTCAGGTGCGAGA-3’, ND3-F:
5’-CCGCGTCCCTTTCTCCATAA-3’, ND3-R:
5’-GGCCAGACTTAGGGCTAGGA-3’, ND4L-F:
5’-TCGCTCACACCTCATATCCTC-3’, ND4L-R:
5’-AGGCGGCAAAGACTAGTATGG-3’, ND4-F:
5’-TTTCCTCCGACCCCCTAACA-3’, ND4-R:
5’-CGTAGGCAGATGGAGCTTGT-3’, ND5-F:
5’-GCCCAATTAGGTCTCCACCC-3’, ND5-R:
5’-GCAGGAATGCTAGGTGTGGT-3’, ND6-F:
5’-ACCCACAGCACCAATCCTAC-3’, ND6-R:
5’-GATTGTTAGCGGTGTGGTCG-3’, COX1-F:
5’-CTTTTCACCGTAGGTGGCCT-3’, COX1-R:
5’-GGCGTAGGTTTGGTCTAGGG-3’, COX2-F:
5’-GCTGTCCCCACATTAGGCTT-3’, COX2-R:
5’-GCTCTAGAGGGGGTAGAGGG-3’, COX3-F:
5’-AGGCATCACCCCGCTAAATC-3’, COX3-R:
5’-CCGTAGATGCCGTCGGAAAT-3’, CYTB-F:
5’-TCTTGCACGAAACGGGATCA-3’, CYTB-R:
5’-GTGGGGAGGGGTGTTTAAGG-3’. The PCR products were then subjected to gel electrophoresis analysis.
.

### Gene knock-out using CRISPR/Cas9

For generation of the CRISPR/Cas9 knockout cell line, gRNA-expressing plasmids were constructed using the px459 vector. Cells were transiently transfected with the appropriate plasmid by Lipofectamine 3000 reagent (Invitrogen) according to the manufacturer’s instructions and screened in the presence of puromycin (2.5 μg/mL for Hela cells and 3.5 μg /mL for 4T1 cells). Single knockout clones were verified by Western Blot and sequencing of the PCR fragment. The following gRNA sequence was used for Hela cells LDHA/B knockout: LDHA-F: 5’-CACCGTCATCGAAGACAAATTGAA-3’, LDHA-R: 5’-CACCGTCATCGAAGACAAATTGAA-3’, LDHB-F:5’- CACCACTTGCTCTTGTGGATGTTT-3’, LDHB-R: 5’- AAACAAACATCCACAAGAGCAAGT-3’. The following gRNA sequence was used for 4T1 cells LDHA/B knockout: LDHA-F: 5’-CACCGCTGGTCATTATCACCGCGG-3’, LDHA-R: 5’- AAACCCGCGGTGATAATGACCAGC-3’, LDHB-F: 5’- CACCGACGGCAGGAGTCCGCCAGC-3’, LDHB-R: 5’- AAACGCTGGCGGACTCCTGCCGTC-3’.

### Treatment of cancer cells with PL, PEITC, DOX and FX11

For cells treated with PL and PEITC, cells were cultured in medium containing 10 μM of PL or PEITC, as described(***Raj, et al.,2011; Trachootham, et al.,2006***). For cells treated with DOX, cells were cultured in medium containing DOX (10 μg/ml). For cells treated with FX11, cells were cultured in medium containing 10 μM of FX11.

### Microscopy imaging of ROS

ROS measurement is assayed by dichloro-dihydro-fluorescein diacetate (DCFH-DA, Sigma), MitoSOX™ Red mitochondrial superoxide indicator (Invitrogen), according to manufacturers’ instruction. Briefly, cells were loaded with DCFH-DA (final concentration of 10 μM for 30 minutes), MitoSOX™ Red (final concentration of 2.5 μM for 10 minutes), washed with ice cold HBSS (Hank’s Balanced Salt Solution, pH 7.2), then observed under a Zeiss LSM710 laser confocal microscope (Carl Zeiss, Germany) equipped with Zen software to process the image. The intensity of fluorescence was analyzed by ImageJ software.

### siRNA knock down of LDH

The following siRNAs targeting LDH were synthesized LDH-A: 5’-GGCAAAGACTATAATGTAA-3’, LDH-B: 5’-GGGAAAGTCTCTGGCTGAT-3’, NC: 5’-GATCATACGTGCGATCAGA-3’. The siRNA was transfer to HeLa cells using RNAiMAX reagent (Thermofisher Scientific, Inc.) according to manufacturer’s protocol. Briefly, after HeLa cell reached 80% confluence in 6-well plate, the siRNA and the RNAi reagent were mixed with Opti-MEM medium and added to plate drop by drop. The cells were subjected to fluorescent microscopy analysis for ROS or collected for LDH activity or western blot analysis 96 hours after transfection.

### Clone and expression of LDH proteins

Total mRNA was extracted from HeLa cell using RNeasy Mini Kit (Qiagen) according to manufacturer’s protocol. The LDHA and LDHB genes were firstly amplified with first round of PCR from total mRNA, then the second round of PCR was used to amplify the coding domain sequence of enzymes with the following primers: LDH-A-1^st^-F: 5’-AGCTGTTCCACTTAAGGCCC-3’, LDH-A-1^st^-R: 5’-GGGTTGCCCAAGAATAGCCT-3’, LDH-A-2^nd^-F: 5’-GGAATTCCATATGGCAACTCTAAAGGATCAG-3’, LDH-A-2^nd^-R:: 5’-ATAAGAATGCGGCCGCATGATATGACATCAGAAGACTT-3’, LDH-B-1^st^-F: 5’- TCCAGAGCCTTCTCTCTCCT-3’, LDH-B-1^st^-R: 5’- GGCTTTGATTCTGTGAGCCC-3’, LDH-B-2^nd^-F: 5’- GGAATTCCATATGGCAACTCTTAAGGAAAAACTC-3’, LDH-B-2^nd^-R: 5’-ATAAGAATGCGGCCGCAGAGCTCACTAGTCACAGGT-3’. The final PCR products were cloned into pET28 a (+) expression plasmid. Then the protein-coding plasmids were transferred into BL21(DE3) *E.coli* by standard transformation protocol. Bacterium pelleted from 250 mL LB culture broth were lysed in PBS with 10 mM PMSF by sonication. After 16000 g and 10 minutes centrifugation, the supernatant was collected for enzyme purification.

The crude LDH proteins were extracted by ammonium sulfate precipitation. Ammonium sulfate was added to protein supernatant slowly with constant stirring on ice. The supernatant from 40% ammonium sulfate solution was precipitated again by adding ammonium sulfate to 60%. After centrifugation, protein pellet was dissolved in PBS with 0.5 M NaCl and 10 mM PMSF. The crude protein solution was kept at 4°C and purified by Ni-Sepharose affinity chromatography later.

The Ni-Sepharose column was first washed and equilibrated by 10 mM imidazole with 0.5 M NaCl (pH8.8). Then the protein solution was loaded into the column. After 20 column volumes’ wash by washing buffer (100 mM imidazole with 0.5 M NaCl, pH8.8), the LDH protein was eluted by 10 column volume of elution buffer (300 mM imidazole with 0.5 M NaCl, pH8.8).

The eluents were concentrated by ultrafiltration using centrifugal filters with cut-off value 10-kD (Millipore).

### LDH enzyme activity assay

Cells maintained in complete growth medium at 70% confluence was washed by PBS and lysed by M-PER buffer (ThermoFisher Scientific). The supernatants after centrifugation (14,000 rpm/30 min/ 4°C) were collected for further analysis. Supernatants or purified enzymes were added to reaction buffer (50 mM Tris-HCl, 1 mM pyruvate, 100 μM NADH, pH 7.4) to initiate reaction. The absorbance at 340 nm wavelength was recorded at 25 °C with a spectrophotometer. For the K_i_ and V_max_ value calculation, FX11 was added to reaction buffer and the initial velocities under difference concentrations of NADH were recorded.

### Measurement of H_2_O_2_-producing activity of LDH

The H_2_O_2_ was detected by Amplex™ Red system. All the reactions were performed in 50mM Tris Buffer with 0.02% BSA, 10μM Amplex™ Red, pH7.4 or other pH value as indicated. Varies substrates, such as NADH, NAD, lactate, pyruvate, were added and the appropriate amount of enzyme was added at last to initiate the reaction. After thorough mixing, 150 μL mixture of each sample was loaded to 96-well black plate. The emission fluorescence at 585 ± 15 nm with the excitation wavelength 525 ± 15 nm was recorded by SpectraMAX i3 Multi-Mode microplate reader (Molecular Devices).

For the inhibition assay, the FX11 and oxamate were added to the reaction mixture at the indicated concentration. Bovine LDH from heart (LDHB) and from muscle (LDHA) were purchased from Sigma-Aldrich.

### Oxygen consumption rate analysis

Cells maintained in complete growth medium at 70% confluence were trypsinized and resuspended in cold fresh medium at the density of 10^6^ cells/mL. Oxygen consumption rate was analyzed on Oxygraph-2K platform (Oroboros Instruments Corp, Austria) according to manufacturer’s protocol. The OXPHOS inhibitors oligomycin (1 μM), FCCP (1 μM), rotenone (1 μM), antimycin (5 μM) were sequentially injected into the assay chambers when the oxygen consumption rate was stable.

### Statistics

All data were analyzed using the GraphPad Prism software (GraphPad Prism 7.0). Two-tailed Student’s t-test was used for statistical analysis, and significance was defined at p<0.05, *; p<0.01, **; p<0.001, ***.

## Acknowledgement

This work has been supported in part by the China National 973 project (2013CB911303), China Natural Sciences Foundation projects (81470126), a key project (2018C03009) funded by Zhejiang Provincial Department of Sciences and Technologies, and the Fundamental Research Funds for the Central Universities, National Ministry of Education, China, to XH, and Zhejiang Provincial Natural Science Foundation of China (LY17H160036), the Fundamental Research Funds for the Central Universities and China Natural Sciences Foundation project 81301707, to HW..

## Author contribution

XH conceived the project, designed the study, XH, HW wrote the paper; HW, YW, and MY performed experiments. XH, HW, YW, MY analyzed data.

---

**Supplementary figure 1.**
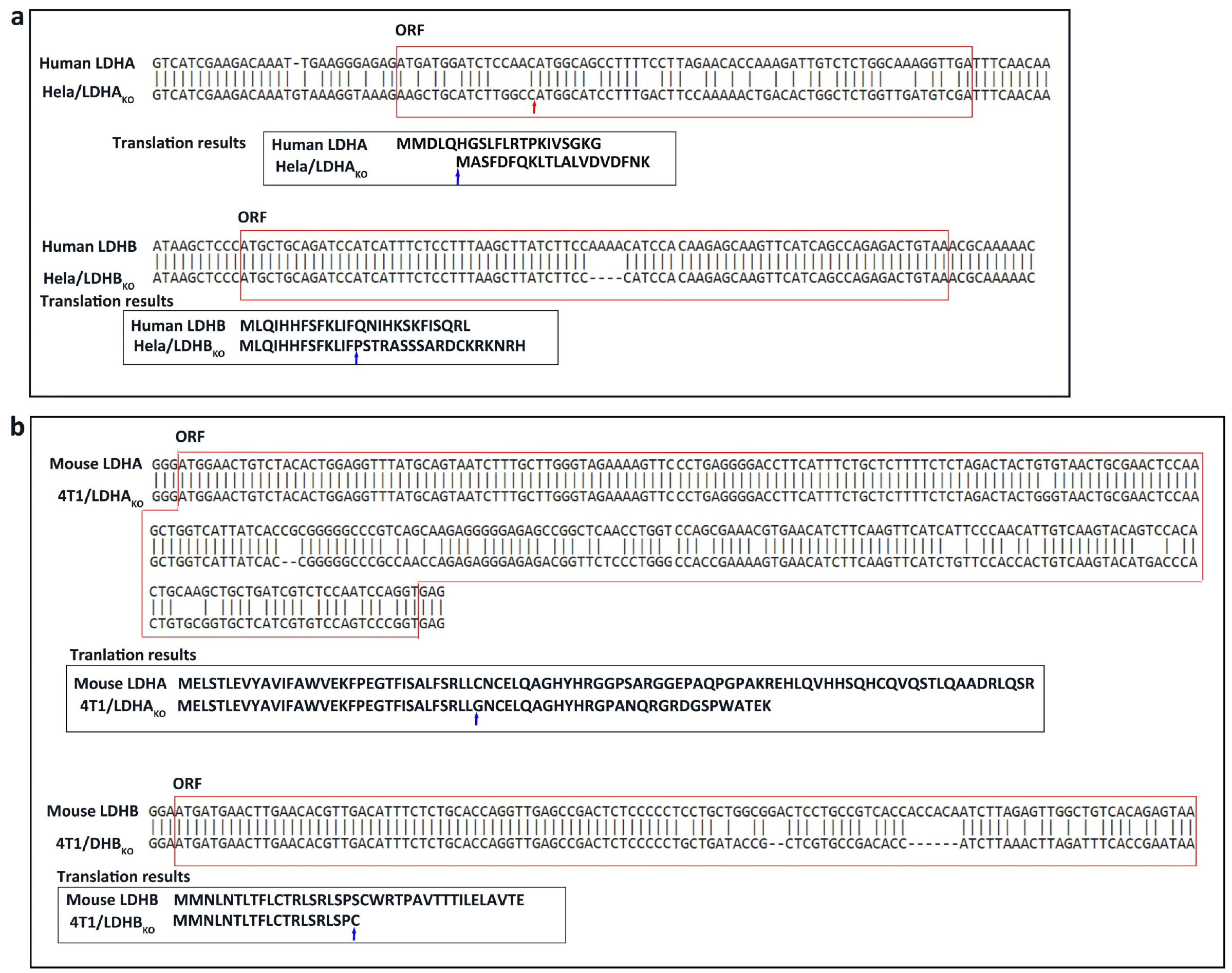
The mutation spectra of target gene. The original gene sequences and each mutant clone were showed in alignment. The red frames showed ORF (Open Reading Frame, ORF) around the target sequences. Between original sequences and mutant clones, the different bases mean point mutant and short lines mean base deletion or insert. The translation results showed protein sequences translated from the ORF respectively. The red arrow showed the altered start codon location of modified genes and the blue arrows showed the first amino acid that different from original peptide. (a) Hela/LDHA_KO_ and Hela/LDHB_KO_ clone cells aligned with human-LDHA, -LDHB original sequences for ORF gene and proteins. (b) 4T1/LDHA_KO_ and 4T1/LDHB_KO_ clone cells aligned with mouse-LDHA, -LDHB original sequences for ORF gene and proteins.

**Supplementary figure 2.**
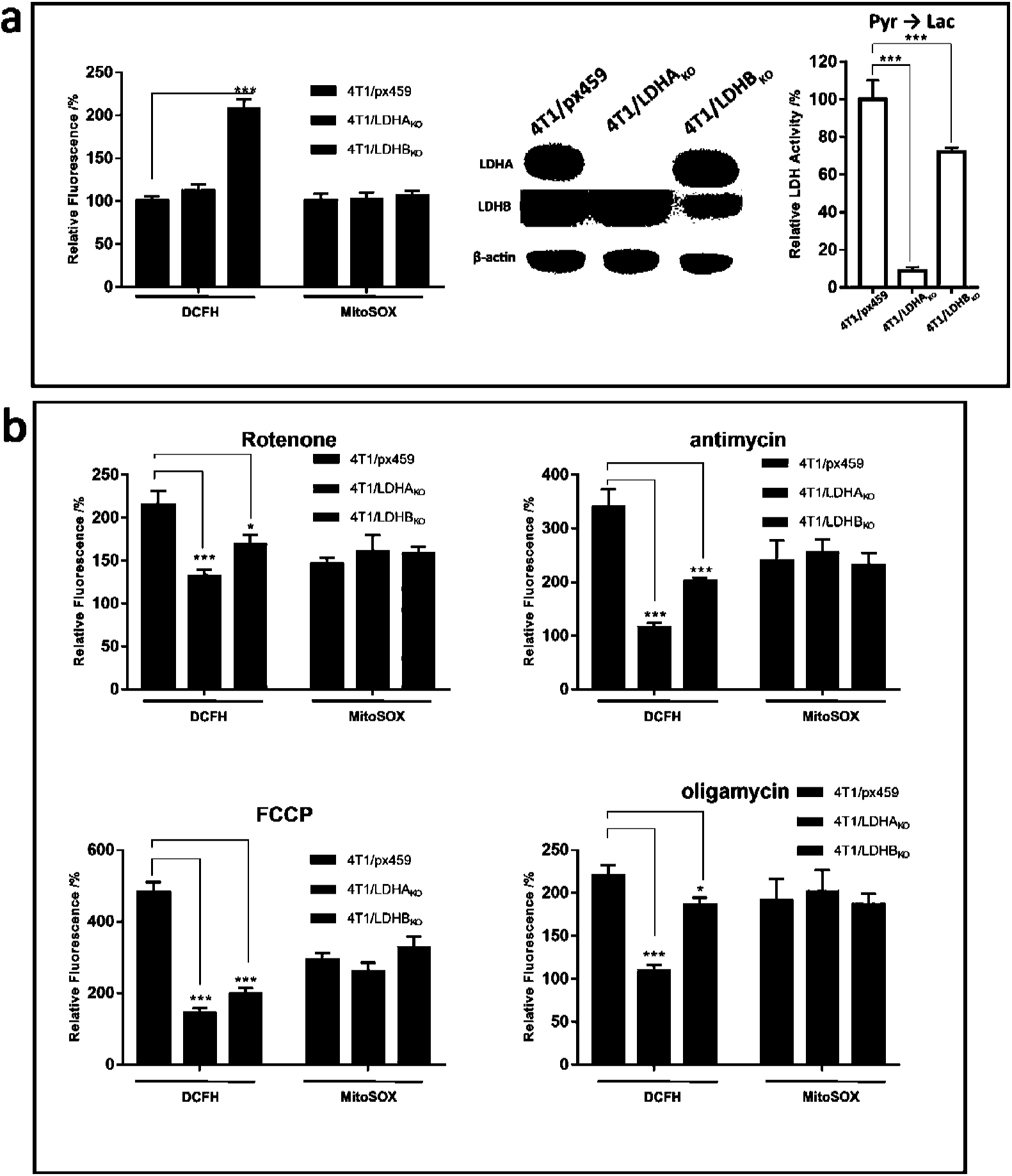
The relationship between cellular ROS and LDH knockout. (a) Characterizations of 4T1/LDHA_KO_, and 4T1/LDHB_KO_ cells. From left to right: relative DCFH and MitoSOX™ Red signals in 4T1/px459, 4T1/LDHA_KO_, and 4T1/LDHB_KO_ cells; LDH activity assayed by converting pyruvate to lactate; Western blot of LDH. (b) rotenone-, antimycin-, FCCP-, or oligomycin-induced production of mitochondrial superoxide (MitoSOX™ Red) and total cellular ROS (DCFH). The fluorescence intensity is relative to untreated Hela/px459 cells, which is 100%. Data were confirmed by at least 2 independent experiments. For experimental details, see Materials and Methods.

**Supplementary figure 3.**
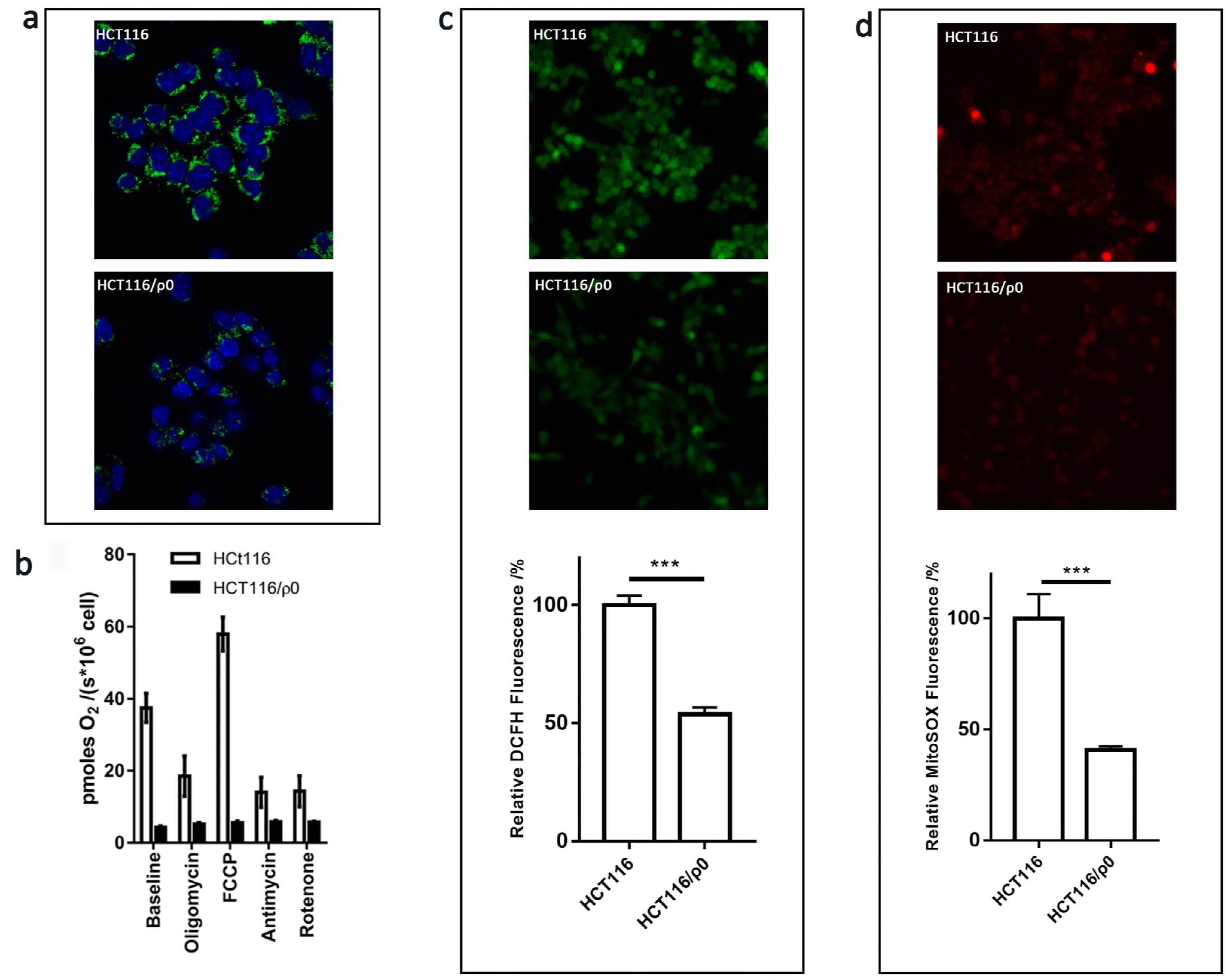
Characterization of HCT116/ρ0 cells. (a) MitoTraker staining of HCT116 and HCT116/ρ0 cells. (b) Oxygen consumption rate in comparison to HCT116 cells; (c) DCFH signal in comparison to HCT116 cells; (d) MitoSOX™ Red signal in comparison to HCT116 cells. Data were confirmed by at least 2 independent experiments. For experimental details, see Materials and Methods.

